# Highly selective visual receptive fields in mouse frontal cortex

**DOI:** 10.1101/2025.03.25.645272

**Authors:** Anthony D. Lien, Bilal Haider

## Abstract

A hallmark of the mammalian visual system is spatial information processing. This relies on feedforward activity spanning multiple brain areas, and on interconnected neurons with spatial receptive fields (RFs) aligned across these areas. This organization allows neurons to iteratively analyze information from the same point of the visual field. It remains unclear if this framework extends beyond the visual system, especially into cognitive areas of frontal cortex that exert feedback control over early sensory areas. Here, we surveyed the mouse frontal cortex (anterior cingulate and secondary motor areas), and discovered neurons with low latency, highly localized visual RFs. Just like in visual cortex, responses were often highly selective for bright or dark stimuli. The responses lagged visual cortical areas by only ∼25 ms, and their RFs were comparable in size. Further, the representation of visual space in frontal cortex showed a strong bias for the central (binocular) visual field, but there was no evidence of a topographically organized retinotopic map. Importantly, these visual responses were abolished by optogenetic silencing of visual cortex, revealing a causal role for feedforward hierarchical connectivity that extends visual spatial processing directly into motor and cognitive regions of mouse frontal cortex.

## Introduction

Feedforward hierarchical processing is a central concept of nervous system function. This has gained strong evidence from the mammalian visual system, where visual signals propagate from retina to thalamus to primary visual cortex (V1), then across multiple hierarchically ordered cortical higher visual areas (HVAs)^1-3^. Each area in turn extracts and integrates visual information passed on from the previous stages of processing, resulting in multiple distinct neural representations across areas, each selective for specific visual stimulus properties^4,5^. This model is foundational for understanding information processing in cortex^3,6-10^, and forms the backbone for artificial intelligence systems^11^.

A key feature of the hierarchical model is that interconnected cortical areas analyze the same part of visual space^2^. This idea relies on foundational discoveries of neurons with spatial receptive fields (RFs) throughout the visual system. The spatial RF corresponds to a discrete region of space where visual stimuli (either bright or dark) drive spiking^12,13^. In V1 of all mammals, neighboring neurons share overlapping RF locations. In this way, neural populations form an orderly topographic map of visual space. Both of these properties (spatial RFs and topography) are also found throughout multiple HVAs, suggesting that hierarchical visual processing maintains spatial alignment across multiple brain regions^2^, even as RF properties such as size, timing, and selectivity for luminance (brights and darks) undergo elaboration in each area^4,5,14^.

There remain several open questions about spatial representations in the feedforward model. First, it is not fully established that spatial RFs faithfully extend “point by point” all the way to higher cognitive areas such as those found in frontal cortex. It is important to understand spatial representations in these areas because they are thought to exert “top down” control over visual functions in HVAs that themselves contain spatial representations^15-19^. Studies in monkey frontal cortex have found neurons with coarse spatial RFs^20-31^, and some evidence for topographic organization of RF locations^21,27,28^, but this can vary widely across studies and experiments that often involve learning and performing complex visual behavioral tasks; further, most of these studies primarily examine the central (foveal) visual field. Second, it is not fully established that visual responses in frontal cortex depend on direct feedforward cortical input from HVAs^32^. This is important to determine because cortico-cortical communication is a key assumption of the feedforward hierarchical model, but subcortical inputs could also directly drive frontal cortex visual responses.^33,34^ Therefore, it would be beneficial to 1) comprehensively map the neural representation of both central and peripheral visual space in frontal cortex, 2) determine the spatial resolution and selectivity of neural representations, and 3) determine that this is an intrinsic property of frontal cortex that depends on direct feedforward cortical input, regardless of learning or visual tasks.

These open questions are now approachable in the mouse. Like primates, the mouse visual system consists of multiple hierarchical and topographically organized HVAs^35-37^, wherein neurons show spatially localized RFs^35,37,38^ and distinct tuning properties ^38-41^. The validity of the feedforward hierarchical model in mice, coupled with the availability of advanced techniques, has propelled detailed insight into the anatomical, cellular, and synaptic basis of visual information processing ^42-48^. However, compared to V1 and HVAs, the functional organization of visual processing in mouse frontal cortex is far less known. There is clear feedforward anatomical input to frontal cortex from the HVAs ^49-57^ and some evidence for visual responsiveness^50,51,58-65^, but the detailed single-neuron substrates and visual selectivity of these areas is unknown. Establishing the functional organization for visual spatial processing in mouse frontal cortex enables comparison across species, and evaluation of general principles underlying the feedforward hierarchical model.

To address these questions, here we mapped spiking responses to classical RF mapping stimuli (flashed bright and dark bars) with Neuropixels probes across the mouse frontal cortex (anterior cingulate cortex, ACC; secondary motor cortex, M2). Surprisingly, we found ∼10% of neurons showed clear, discrete visual spatial RFs; many of these were also highly selective for bright versus dark stimuli. Frontal cortex RF responses lagged those in HVAs by only ∼25 ms, and their RFs were comparable in size. Frontal cortex RFs were strongly biased to the central (binocular) visual field, and there was no evidence for topographic organization of retinotopic preferences. Lastly, these frontal cortex RFs were abolished during inactivation of V1 or HVAs, establishing that these highly selective visual spatial representations in mouse frontal cortex originate from feedforward visual cortical activity.

## Results

### Highly selective visual receptive fields in mouse frontal cortex

We first set out to determine if neurons in mouse frontal cortex showed spatially specific visual responses. We targeted Neuropixel recordings to the medial region of frontal cortex in awake mice that passively viewed visual stimuli while sitting in a tube (Fig. 1A). Recordings spanned medial secondary motor (M2) and dorsal and ventral anterior cingulate cortices (dACC and vACC) as confirmed by histological reconstruction of probe tracks in a subset of mice (23/47 recordings from 5/14 mice). These regions are known to be reciprocally connected with V1 and higher visual areas ^49-57,66^ (Fig. 1B schematic). We focused our analysis on putative excitatory neurons with regular spiking (RS) waveforms although neurons with fast spiking waveforms (putative inhibitory interneurons) had similar properties (Fig. S1). To map visual spatial receptive fields, we presented single black or white vertical bars against a gray background at different locations spanning ∼140° of horizontal (azimuth) space in both binocular and monocular visual fields (Fig. 1A schematic).

**Figure 1.**
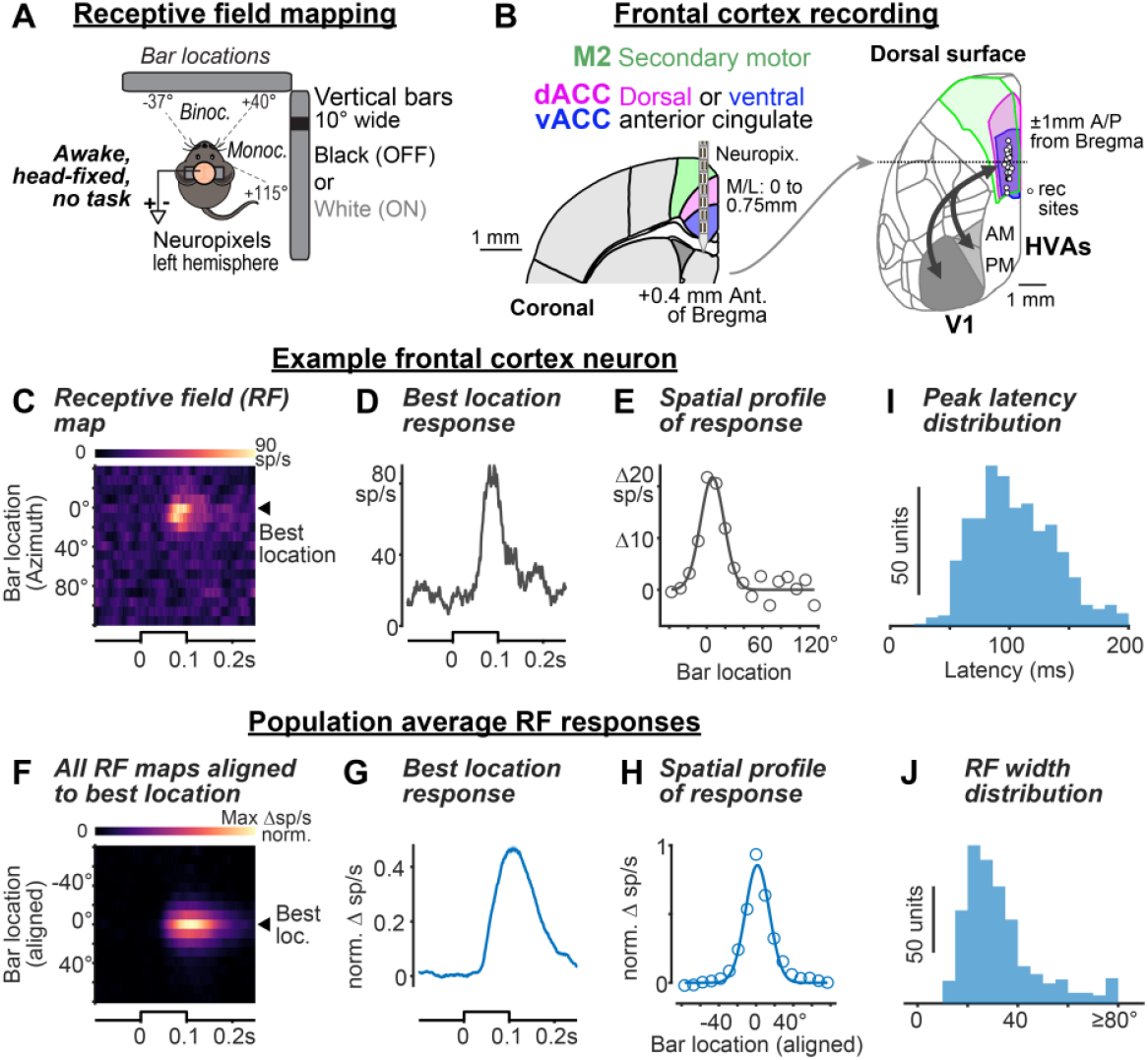
Spatially selective visual receptive fields in frontal cortex. A. Receptive field mapping schematic. Awake head-fixed mice sat in a tube in front of two screens showing vertical bars (one at a time) at different horizontal (azimuth) locations. Vertical meridian straight ahead of mouse defined as 0°. *B. Left*, coronal atlas section (+0.4mm anterior of bregma) showing Neuropixels probe spanning secondary motor (M2) and anterior cingulate cortex (dorsal, dACC; ventral, vACC). *Right*, dorsal surface view of reconstructed frontal recording sites (n=23 recordings; 5 mice), spanning ±1mm A/P from Bregma. Arrows indicate known reciprocal projections between frontal and primary visual cortex (V1) and anteromedial (AM) and posteromedial (PM) higher visual areas (HVAs). C. Receptive field (RF) map of an example frontal cortex regular-spiking (RS) neuron. Triangle indicates best location (best bar position eliciting maximal response). Only responses to white bars shown. D. Same neuron as C, showing time course of best location response. E. Same neuron as C, showing spatial profile of RF responses (circles) with Gaussian fit (solid). F. Population RF map computed after aligning each neuron’s best bar response (color and location). (n=589 RS units, 47 recordings, 14 mice in F-J). G. Population average spiking response to the best bar location. H. Population average spatial profile (circles) with Gaussian fit (solid) I. Peak latency distribution calculated from each unit’s best bar response. (100.5 ± 23.0 ms, median ± MAD). J. Distribution of RF width calculated from each unit’s spatial profile (29.1 ± 7.3°, median ± MAD).

Surprisingly, we found that ∼10% of all recorded neurons in frontal cortex (710/6757) showed clear, spatially-localized visual receptive fields (RFs). Fig. 1C shows the RF map of an example regular spiking (RS, putative excitatory) neuron in response to bars of its preferred color (in this case, white). This neuron responded with short latency visually-evoked spikes (within 0.1s) following the onset of bars presented at -10° to 20° (Fig. 1D). The spatial profile of spiking responses was well-fit by a Gaussian centered at 6.1° with a width of 29.8° (full-width at half-max, Fig. 1E). The average RF map of all frontal RS neurons with quantifiable RFs (Fig. 1F, 589 / 5990 RS neurons, aligned to each neuron’s best bar location) showed a tightly localized spatial response with median width of 29.1 ± 7.3° and peak latency of 0.100 ± 0.023s across the population (Fig. 1G-J). Fast spiking (FS, putative inhibitory) neurons also showed spatially selective RFs (Fig. S1; 16% of all FS units; median width of 36 ± 9º, significantly broader than RS; *p* = 6.97e-7, rank sum test). Further, showing horizontal bars revealed these spatially localized RFs centered just above eye level in vertical space (elevation) (Fig. S2). We will focus the remaining analysis on selectivity for horizontal visual space in RS neurons, since these are the majority of cortical neurons and the main source of frontal cortical output^54,67,68^.

Remarkably, frontal cortex neurons responded selectively to increases (ON) or decreases (OFF) in luminance, just like neurons in the early visual system. We found an unexpected degree of OFF versus ON selectivity, visible in RF maps constructed from either black or white bars. Some neurons responded exclusively to one bar or the other, while others responded equally well to both (Fig. 2A-B). To visualize the strength of OFF/ON selectivity across the population, we first computed a selectivity score for each neuron (Fig. 2C), based on the response magnitude to black versus white bars (Methods), then plotted the average RF responses to black and white bars for neural subpopulations sorted by their selectivity (Fig. 2D). Around 22% of neurons with RFs were strongly selective for white or black bars (13% ON selective, Group 1; 9% OFF selective, Group 5), 34% were equally responsive to white or black bars (Selectivity Group 3), with the remaining neurons exhibiting intermediate degrees of selectivity. Frontal cortex FS neurons were significantly less selective for bar luminance than RS neurons (absolute value of OFF/ON selectivity score RS: 0.3 ± 0.18; FS: 0.25 ± 0.16; *p* = 0.027; median ± MAD, rank sum test), echoing the lower selectivity of FS neurons in visual cortex ^69-72^.

**Figure 2.**
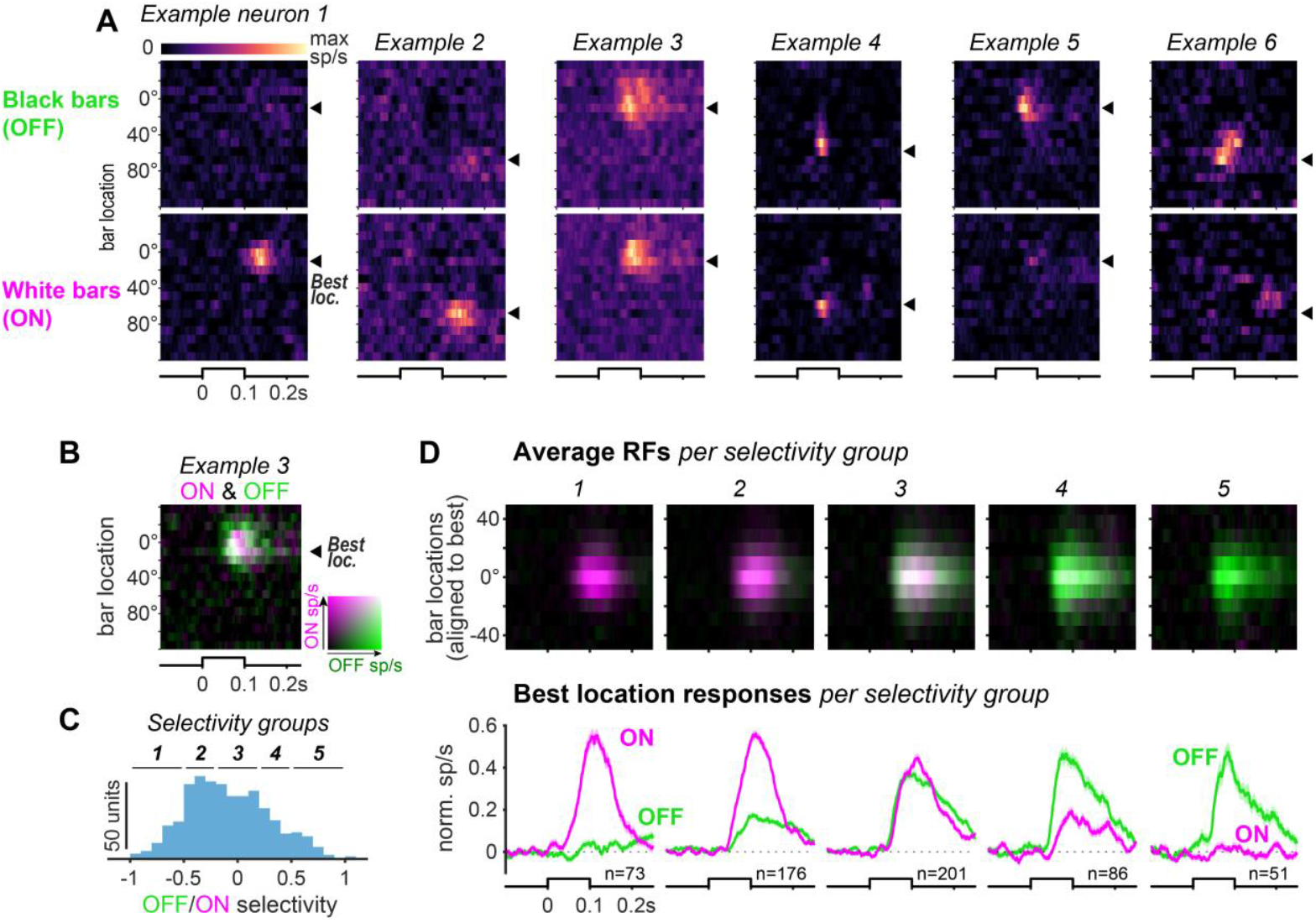
ON/OFF selectivity of frontal cortex receptive fields. A.Six example neuron RF maps for OFF responses to black (top) or ON responses to white (bottom) bars. Arrowhead at location for best response averaged across all bar colors *B*.Merged ON and OFF RF maps map for example neuron 3 from A. *Lower right*, color saturation shows strength of ON (magenta) or OFF (green) response, while mixed selectivity response (diagonal) strengthens from black to white values. C.istribution of OFF/ON selectivity index. Negative values denote preference for white bars (ON), positive values for black bars (OFF). Neurons were divided into 5 selectivity groups (horizontal lines at top; n=589 RS units). D.*Top*, population average of merged and aligned ON (magenta) and OFF (green) RF maps. *Bottom*, population spiking response for black or white bars (average response at center (0°) ± 1 adjacent locations in RF maps at top), for each selectivity group in C

Are all parts of horizontal visual space represented in mouse frontal cortex? Indeed, neurons had RFs that tiled across visual space (example units in Fig. 2A). However, when we plotted the overall population RF maps, it was clear that the population response was more strongly driven by bars in binocular (-40° to 40°) rather than monocular (>40°) visual space (Fig. 3A, right; best binocular response: 5.4 ± 3.8 spikes/s; best monocular response: 1.7 ± 1.6 spikes/s *p* = 4.7e-67; median ± MAD; sign rank test). Overall, 85% of neurons had RFs in the binocular field compared to only 15% in the monocular field (Fig. 3B). The binocular bias of frontal cortex receptive fields was observed in each individual mouse (Fig. S3) as well as in FS neurons (Fig. S1i), indicating that binocular visual space is strongly represented in the mouse frontal cortex.

**Figure 3.**
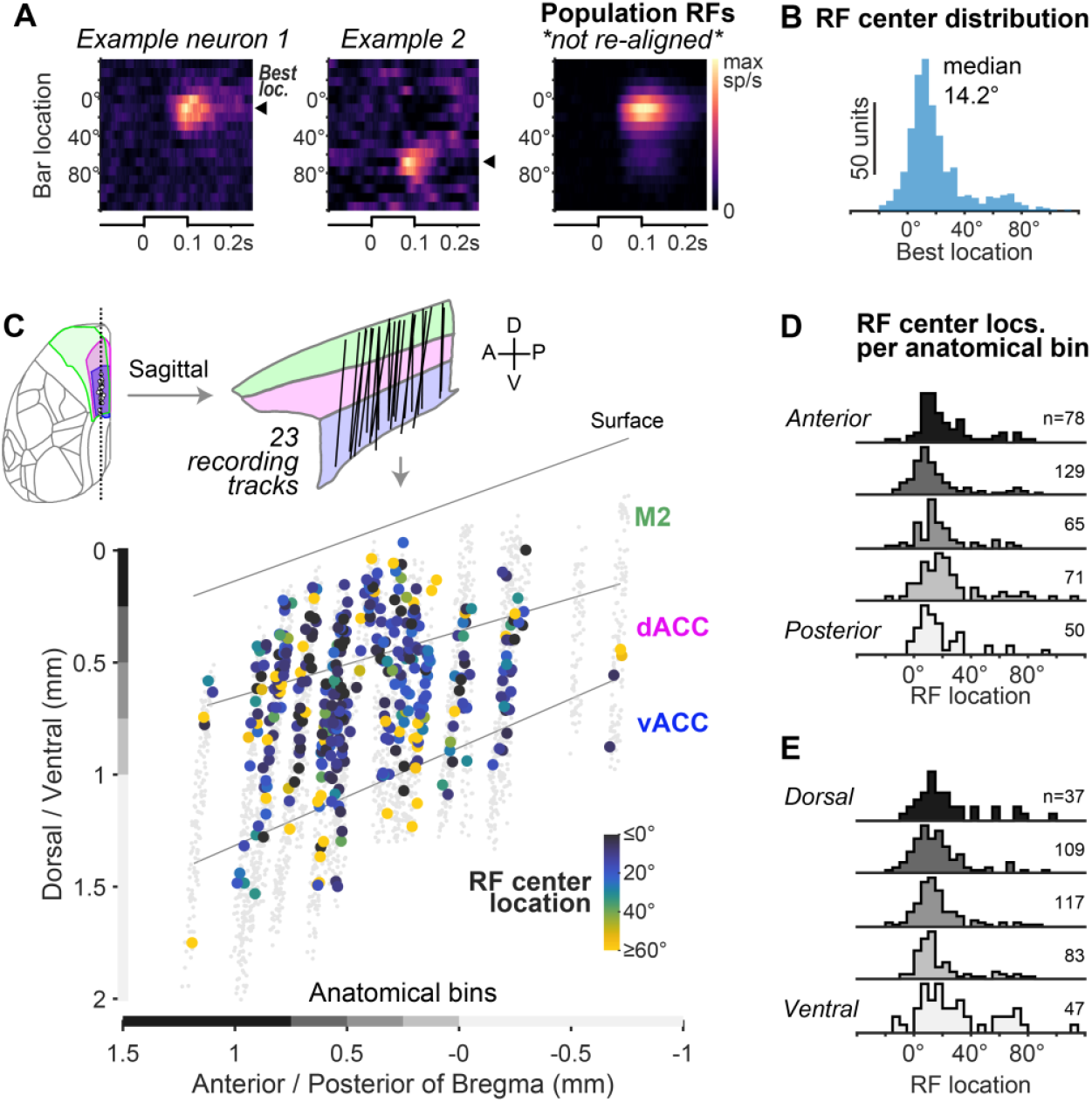
Binocular bias and lack of topographic organization for visual neurons in frontal cortex. A. *Left and middle*, RF maps of frontal cortex neurons with binocular (*Example 1*) or monocular (*Example 2*) selectivity (see Fig. 1A visual field schematic). Triangles (best bar location) denote RF center. *Right*, Population average RF map (without re-alignment of best bar location; n=589 RS neurons, 47 recordings, 14 mice). *B*. Distribution of RF center locations (14.2 ± 8.5°; median ± MAD; same neurons as A). C. *Top*, all recording tracks with fluorescent label (n=23), projected in the sagittal plane. *Bottom*, neurons with significant visual RFs color-coded by center location (n=393 RS neurons), all other non-RF neurons in gray (n=3370 RS neurons). Lines denote area boundaries. Gray and black bars above axes indicate anatomical bins used in D and E (n=23 recordings, 5 mice; same units for D-E). For display purposes, neuron coordinates were randomly jittered by ≤25 μm in each dimension. D. Distribution of RF center locations across the anterior/posterior axis using the 5 anatomical bins depicted in D. No correlation between RF location and anterior/posterior coordinate (r=0.017, *p*=0.73; Spearman correlation). E. Same as D for the dorsal/ventral axis. No correlation between RF location and dorsal/ventral (r=0.066, *p*=0.19; Spearman correlation) or medial/lateral coordinate (r=0.028, *p*=0.58; Spearman correlation).

Across the frontal cortex, we found little evidence for large-scale topographic organization of RF locations (retinotopy). In a subset of experiments, we used histological reconstruction of probe tracks to register the locations of individual neurons to the Allen Brain Common Coordinate Framework^73^ (Methods). Each recorded neuron was plotted at its anterior/posterior and dorsal/ventral coordinates in the sagittal plane and color-coded by RF location (Fig. 3C). The monocular units were intermingled with binocular units and the RF location showed little correlation with either anterior/posterior, dorsal/ventral, or medial/lateral coordinates (Fig. 3D-E). Furthermore, the strong bias for binocular RFs was maintained in anatomical subregions binned across both anterior/posterior and dorsal/ventral dimensions (Fig. 3D-E).

### Comparison of frontal RFs to V1 and higher visual areas (HVAs)

How do temporal and spatial RF properties in frontal neurons compare with those in primary and higher visual cortex? We first compared the temporal properties of RF responses across V1, HVAs, and frontal cortex. We focused on the anteromedial and posteromedial HVAs (AM/PM) because they exhibit particularly high connectivity with frontal cortex^49-53,66^, as we verified in the Allen Brain Connectivity Atlas^55^ (Methods). We recorded from V1 and AM/PM and mapped RFs using the same stimuli used for frontal cortex (Fig. S4A-B).

As expected, V1 and AM/PM showed much higher fractions of visual RF units (V1 28% 258/939, AM/PM 29% 289/1005) than frontal cortex (10% 589/5990). V1 had significantly faster visual response latencies than AM/PM (V1 time to peak 64 ± 43 ms, AM/PM 74 ms, median ± IQR; *p* = 1.7e-4, post hoc Dunn’s test; Fig. S4D-E). Frontal cortex neurons had the longest response latencies, lagging AM/PM latencies by ∼25ms (101 ± 47 ms, median ± IQR; *p* = 4.9e-19, post hoc Dunn’s test; Fig. S4D-E). This timing is consistent with frontal cortex RF neurons being functionally downstream of both V1 and HVAs. We next examined spatial properties of RFs, and found that frontal cortex RF widths were only slightly greater than AM/PM (by ∼2°; *p* = 0.036), while both frontal and AM/PM RFs were ∼12-14° greater than V1 (V1: 17.4 ± 9.2°; AM/PM: 27.0 ± 18.3°, *p* = 2.4e-27; Frontal: 29.1 ± 15.0°, *p* = 2.6e-47; median ± IQR, post hoc Dunn’s test; Fig. S4F-G). Thus, RF neurons in mouse frontal cortex show a similar degree of spatial selectivity as neurons in HVAs.

### Visual cortical origin of frontal cortex receptive fields

We found that frontal neuron visual responses depended on feedforward input from V1 and HVAs ^60^. We mapped RFs of frontal cortical neurons (including both RS and FS neurons) while optogenetically silencing visual cortex (on a fraction of interleaved trials) to compare how frontal neuron RFs changed with and without visual cortical input. We targeted a blue laser spot (36 µm full-width at half max) to the skull surface over binocular V1 (V1B), monocular V1 (V1M), or AM/PM in mice expressing channelrhodopsin (ChR2) in parvalbumin (PV) inhibitory interneurons to silence visual cortical activity ^48,74,75^ (Fig. 4A). Recordings at the site of optogenetic silencing showed activation of FS neurons and complete silencing of RS neurons during receptive field mapping (Fig. S5A-B). Optogenetic silencing of each of the three visual cortical sites significantly reduced RF responses in frontal cortex (Fig. 4B-E). During silencing of V1B, the response of frontal cortex neurons to bars across the entire RF was almost completely abolished compared to interleaved control trials (Fig. 4B-E, top). Further, the amount of response reduction was significantly greater when targeting V1B compared to V1M and AM/PM (Fig. 4E). With either V1M or AM/PM silencing, frontal cortex neurons showed significant response reductions, but these were more gradual across the RF extent (Fig. 4C), and caused comparable reduction of best bar responses (Fig. 4E; V1M vs AM/PM, *p* = 0.26, post hoc Dunn’s test). These results demonstrate that the visual receptive fields in frontal cortex depend on feedforward visual cortical activity, with binocular V1 having the largest impact.

**Figure 4.**
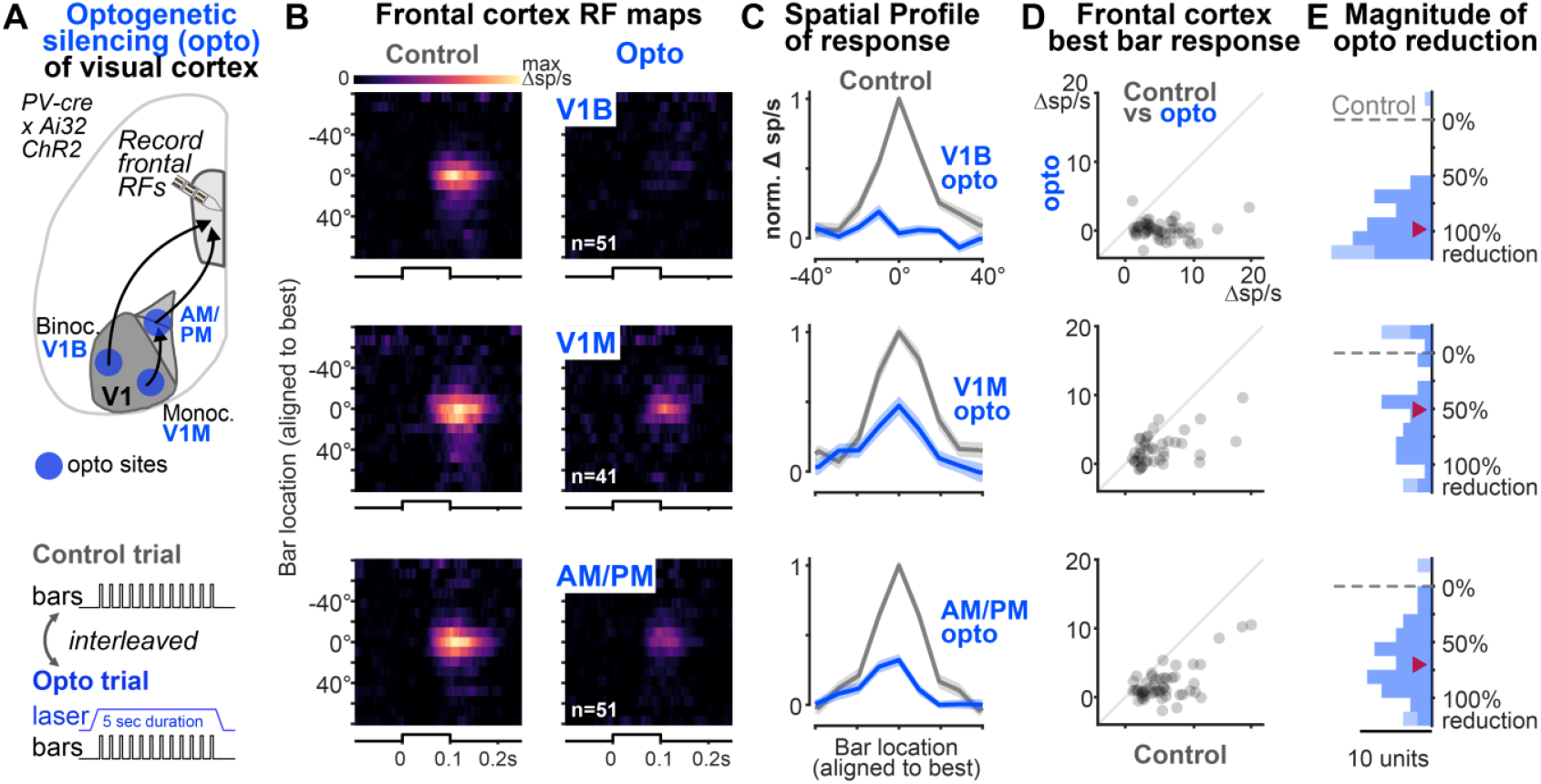
Visual neurons in frontal cortex require input from visual cortex. A. *Top, e*xperimental approach for optogenetic silencing (“opto”) of visual cortex while mapping frontal cortex RFs. Laser targeted three separate sites (V1M, monocular V1; V1B, binocular V1; AM/PM border). Sites targeted via ISI maps (Fig. S4B). *Bottom*, neural responses to bars measured during interleaved control and opto trials. *B*. Frontal cortex RF maps in control (left) and opto (right) conditions, with laser targeted to V1B (top row B-E; n=51 frontal neurons, 6 recordings, 2 mice), V1M (middle row B-E; n=41 neurons, 4 recordings, 2 mice), or AM/PM (bottom row B-E; n=51 neurons, 7 recordings, 2 mice). Maps aligned at 0º to each neuron’s best bar location before averaging. Best location determined from control RF mapping prior to any optogenetic experiment. C. Same data as B, showing average spatial profiles of frontal cortex RF neurons in control (gray) and opto (blue) trials for each opto site (rows). Data aligned at 0º to each unit’s best bar location. D. Same data as B and C, showing each frontal neuron’s visually-evoked response magnitude during control (x-axis) and opto (y-axis) trials at each visual cortex site. Responses to each unit’s best bar (0º in C). Line indicates unity. All sites caused significant reduction of peak RF responses (V1B control: 4.7 ± 2.7 spikes/s; opto: 0.1 ± 1.0; *p*=1.6e-9; V1M control: 3.3 ± 2.8 spikes/s; opto: 1.7 ± 1.7; *p*=7.6e-6; AM/PM control: 5.1 ± 2.8 spikes/s; opto: 1.0 ± 1.8; *p*=1.6e-9; Median ± MAD; sign rank test). E. Distribution of frontal neuron response reduction for each opto site in visual cortex. Response reduction was calculated as the percent decrease of baseline-subtracted responses to each unit’s best bar (0º in C) in opto trials relative to control trials. Values > 100% indicate that opto reduced responses below baseline; values < 0% indicate response enhancement. Lighter blue bars contain units with values outside of the plotted range (-25 to 125% reduction). Triangle indicates median. Opto over V1B caused a significantly greater reduction than opto over V1M or AM/PM. (V1B: 98.2 ± 18.7%; V1M: V1M: 50.8 ± 35.0%; AM/PM: 70.2 ± 24.2%; Median ± MAD; *p* = 1.4e-7 Kruskal-Wallis test; V1B vs V1M: *p* = 5.7e-7; V1B vs AM/PM: *p* = 3.1e-5; V1M vs AM/PM *p* = 0.26; post hoc Dunn’s test).

## Discussion

Our study establishes the functional organization of visual selectivity in the mouse frontal cortex. We found a distinct representation of visual space composed of neurons with fast spatial RF responses that were selective for both luminance increases and decreases. Visual representation showed a strong bias for the binocular visual field, but it was not topographically organized into a retinotopic map. Importantly, these frontal RF responses were eliminated by optogenetic silencing of visual cortex, establishing a causal role for feedforward hierarchical cortical processing that extends all the way into the frontal cortex of the mouse.

### Multiple aspects of visual selectivity in mouse frontal cortex

We show that three key forms of visual selectivity are intrinsic properties of frontal neurons. First, they show strong spatial selectivity, evidenced by clearly localized RFs. Previous studies of mouse frontal cortex (showing large stimuli in just 2 locations in the visual field) found coarse selectivity for contralateral stimuli ^50,59,60,64,65^, but these responses provided no hints that classical RF mapping stimuli could drive robust responses that allow detailed quantitative insights about RF structure, as we show here. Surprisingly, RF widths of frontal neurons were comparable to those in medial HVAs, and only ∼70% broader than V1. Thus, frontal neurons show RFs that are nearly as spatially selective as V1 neurons, despite being in premotor and cognitive areas on the opposite end of the brain.

Second, frontal neurons show a high degree of selectivity for luminance increases or decreases (ON/OFF selectivity). ON and OFF pathways are segregated throughout the early visual system (retina, LGN) but heavily converge at the level of V1 (i.e., simple and complex cells)^76^. It is thus unexpected to find such strong ON/OFF selectivity preserved in some frontal neurons. This reveals that they not only encode specific spatial locations, but also selectively encode relative brightening or darkening of the visual field at these specific locations.

Third, the RF locations were strongly biased for the binocular visual field. This is unexpected given that mice lack a fovea, and have a relatively small binocular visual field (due to the lateral position of the eyes) ^77^. The strong binocular bias in frontal cortex stands in stark opposition to V1 and HVAs, which devote most of their territory to neurons with RFs in the monocular visual field, although some HVAs do show relatively greater binocular field coverage^36,78,79^. Thus, frontal cortex appears to share a principle of higher visual cortex by biasing processing to a particular region of the visual field.

Despite these multiple dimensions of visual selectivity in single neurons, there was little evidence for a topographically organized retinotopic map in frontal cortex. This is a major difference from mouse V1 and HVAs, which all show topographic organization of retinotopic preferences ^36,39^. Instead, frontal cortex RFs were organized in a “salt-and-pepper” fashion, with a small population of monocular RFs distributed throughout a sea of binocular RFs. This arrangement is similar to the organization of orientation preferences in rodent V1, where sharp single-neuron orientation tuning is maintained despite disparate tuning preferences in the nearby neurons^80^.

### Frontal cortex visual representations arise from feedforward cortical input

Optogenetic silencing of V1 and HVAs abolished visual responses in frontal cortex neurons. This strongly suggests a visual cortical origin for visual selectivity in frontal neurons, rather than major driving contributions from thalamus (e.g., tecto-thalamo-frontal pathways)^33,34,49,81-85^. However, our experiments cannot distinguish the relative contributions of direct visual cortical projections versus alternative indirect pathways (such as cortico-thalamo-cortical circuits linking visual and frontal cortices^49,81^). Our inactivation results are consistent with a prior study showing reduced bulk Ca^2+^ fluorescence in frontal cortex after pharmacological silencing of visual cortex^60^. Furthermore, we found that frontal cortex visual responses were most dependent on input from binocular V1, matching the binocular bias for RF locations in frontal cortex. This shows retinotopic alignment in the feedforward information from visual to frontal areas. Silencing HVAs AM and PM also reduced frontal cortex RF responses, but to a lesser degree than silencing V1. This could be because of several reasons, including mismatched retinotopic preferences (despite strong anatomical connectivity) between frontal cortex and AM/PM, much stronger contribution from direct V1 inputs, or due to V1 inactivation also reducing HVA activity. This last possibility of cascading effects is consistent with our findings of sequential activation of V1, HVAs, then frontal cortex. However, our control measures in AM/PM showed modest activity reduction during V1 silencing (Fig. S4E); further, we did not record from or attempt to silence other HVAs, particularly those with binocular biases (LM, RL). Future experiments that specifically disrupt synaptic release from V1 and HVA axons in frontal cortex^86^ could further dissociate the contributions of V1 and HVAs to frontal visual responses.

### Implications for mechanisms underlying frontal cortex visual selectivity

Our findings suggest several possibilities for circuit mechanisms driving visual selectivity in frontal cortex. The relatively small increase in RF size from V1 and HVAs to frontal cortex implies low RF scatter and high convergence of the presynaptic visual inputs to individual frontal neurons. Similarly, the ON/OFF selectivity of individual frontal neurons implies a highly specific functional convergence of ON-inputs and OFF-inputs that are also retinotopically aligned. Our results also suggest a role for local excitatory and inhibitory visual processing within frontal cortex^87^. We found that FS putative inhibitory interneurons also have spatially localized RFs, consistent with prior reports of their visual responsiveness and drive from visual cortical excitatory inputs^51,54,83^. Detailed RF mapping showed that frontal FS neurons are less selective than putative excitatory RS neurons, with broader RFs and more mixed ON/OFF selectivity, echoing findings in visual cortex^69-72^. The tight spatial and temporal visual spiking of RS cells implies that they also preserve highly aligned “like to like” local excitatory connectivity^74,88,89^. This must particularly be the case for neurons with monocular RFs which are surrounded by neurons with binocular RFs, again implying highly specific local recurrent connectivity in both excitatory and inhibitory subnetworks^90,91^. Thus, frontal and visual cortices may utilize similar principles of feedforward and local circuit connectivity to generate visual selectivity. These findings raise exciting possibilities for detailed studies of synaptic connectivity in frontal visual neurons using a wide variety of techniques available in mice^54,89,92,93^.

### Implications for feedback from frontal to visual cortex

It is believed that frontal cortex feedback to visual areas enables top-down control of information processing, as relevant for many behaviors requiring selective visual processing^17,30,94^. Much work in primates^15,16,95-98^ suggests this feedback relies on broad “point to point” retinotopically aligned connectivity between neurons sensitive to the same region of visual space. Our findings carry new implications for this scheme in mice. First, frontal cortex responses are rapid and overlap in time with responses in V1 and HVAs, suggesting that feedback modulation can readily occur at a timescale relevant for fast visual processing during behavior. Indeed, frontal feedback circuits in mice recruit monosynaptic excitation in V1 excitatory neurons and multiple inhibitory neuron classes^67,68,99^. Second, only subsets of frontal neurons show localized RFs, begging the question if these neurons provide the major source for feedback projections; alternatively, the visual RF neurons may also recruit “non visual” neurons in frontal cortex to distribute feedback operations across larger populations^50,61,100,101^. Since there is no topographic map of visual space in frontal cortex, retinotopically-matched feedback must rely on fine-grained retinotopic projection specificity at the single neuron level in downstream targets. Identifying the projection targets of frontal cortex neurons with known RF locations, the layers and cell types they target, and their impact on visual cortical activity will be important for understanding the circuit mechanisms of frontal feedback to visual cortex.

Finally, our findings suggest that the binocular visual field is particularly important for mouse frontal cortex. This region of visual space shows specialized processing throughout the mouse visual system^36,78,102,103^. Behaviorally, the binocular region shows greatest perceptual sensitivity and spatial acuity^78,104^. Mice use head and eye movements to maintain binocular fusion for both eyes just above and in front of the nose, a critical factor for successful visual capture of prey during hunting^105-108^. Our findings show that binocular prioritization extends to the frontal cortex, even in the absence of learning, rewards, or task engagement; moreover, our findings highlight that stimulus properties such as location, size, and luminance will be important for understanding the role of mouse frontal cortex in visual behavior. How these frontal neurons contribute to behaviors that require top-down visual feedback, such as during spatial attention^104,109-113^ forms an exciting topic for future work that is now amenable in mice and could provide circuit-level detail for seminal ideas about top-down processing^114^. Our discovery here of visual RFs in multiple regions of frontal cortex puts vision at the front of the mouse brain, and opens the door to dissect the functional architecture of these neurons using a rigorous framework that has proved immensely powerful for understanding visual functions throughout the brain^39,40,48,69,74,115-122^.

## Acknowledgements

We thank members of the Haider lab for feedback, and Aman Saleem for comments on the manuscript. This work was funded by the National Institute of Neurological Disorders and Stroke and NIH BRAIN Initiative (grant nos. NS107968, NS109978, NS132288 to B.H.) and the Simons Foundation (grant no. SFARI 878393, B.H.).

## Author Contributions

A.D.L. performed all experiments and analysis. A.D.L. and B.H. wrote the paper together.

## Lead contact

Further information and requests for resources should be directed to and will be fulfilled by the lead contact, Bilal Haider.

## Materials availability

This study did not generate new unique reagents.

## Data and code availability

All data structures and code that generated each figure will be publicly available at DOI 10.6084/m9.figshare.28646108 upon publication and linked from the corresponding author’s institutional webpage upon publication.

## Methods

### Experimental subject details

C57BL/6J (RRID: IMSR_JAX:000664) mice (referred to as WT), Emx1-Cre (IMSR_JAX:005628) mice crossed with WT mice (referred to as Emx1-Cre), and B6 Pvalb-IRES-Cre (RRID: IMSR_JAX:017320) mice crossed with Ai32(RCL-ChR2(H134R)/EYFP) (RRID: IMSR_JAX:024109) mice (referred to as PV-ChR2) were bred in house. After headplate implantation, mice were individually housed under reverse light cycle conditions.

| Mouse strain | n=mice | n=total frontal cortex recordings (mice) | n=frontal cortex recordings without histology (mice) | n=frontal cortex recordings with histology (mice) | n=V1 recordings (mice) | n=AM/PM recordings (mice) | n=frontal cortex recordings with opto (mice) | n=visual cortex recordings with opto (mice) |
| --- | --- | --- | --- | --- | --- | --- | --- | --- |
| WT | 7 (4 female) | 27 (7) | 7 (3) | 20 (4) | 8 (2) | 8 (2) | 0 | 0 |
| Emx1-Cre | 4 (1 female) | 11 (4) | 8 (3) | 3 (1) | 2 (1) | 7 (2) | 0 | 0 |
| PV-ChR2 | 3 (2 female) | 9 (3) | 9 (3) | 0 | 4 (1) | 2 (2) | 7 (2) | 2 (1) |

All experimental procedures were approved by the Institutional Animal Care and Use Committee (IACUC) at the Georgia Institute of Technology.

### Implant surgery

Mice were implanted with a stainless steel headplate with recording chamber (8mm inner diameter) under isoflurane anesthesia (3% induction, 1-2% maintenance). The frontal cortex site for future craniotomy was marked with a pencil (-0.5 to +0.5 mm AP, 0 to +0.5 mm ML from Bregma) and the headplate was adhered to the skull with veterinary adhesive (VetBond) followed by clear dental cement (Metabond). The skull surface within the recording chamber was covered with a thin layer of cyanoacrylate glue (Zap-a-gap) to maintain optical clarity for intrinsic signal imaging over the left visual cortex. The recording chamber was sealed with silicone elastomer (KwikCast). Mice were given analgesia (ketoprofen 5 mg/kg SC and buprenorphine sustained release 1 mg/kg IP) and allowed to recover for at least 3 days after implantation. In a subset of mice, adeno-associated virus (AAV) for expressing opsins were injected in frontal cortex for optogenetic experiments that were part of separate studies.

### Intrinsic signal imaging

V1 and HVAs were targeted for recording and optogenetics using transcranial intrinsic signal imaging of visual cortex retinotopy as previously described ^123,124^. Mice were anesthetized and head-fixed under light isoflurane anesthesia (0.7-1%) and sedation with chlorprothixene (1 mg/kg IP). Contact lenses were used to maintain optical clarity of the eye. Mice were positioned under the imaging system and reference images of brain vasculature in the left visual cortex were captured under green light (475-590 nm) illumination. Hemodynamic responses were acquired at 10 frames per second under red light (>610 nm) illumination at a focal plane ∼0.5 mm beneath the brain surface during visual stimuli that drifted periodically across two monitors covering 150° (horizontal) by 48° (vertical) of the visual field. Azimuth (horizontal) and elevation (vertical) retinotopic maps were constructed from phase maps of the hemodynamic response. Visual field sign maps were calculated as the sine of the difference in gradient angle of azimuth and elevation maps for each pixel. Area AM was identified as the negative field sign patch at the anterior side of the medial border with V1 and PM the positive field sign patch immediately posterior of AM (see Fig. S4B).

### Visual stimuli

Visual stimuli were presented to head-fixed mice on a pair of LCD monitors positioned in front of and to the right of the animal (Fig. 1A). The luminance of white, gray, and black values was 190, 100, and 5 cd/m^2^ in most experiments, except a few where peak positive luminance values were 300 cd/m^2^. These differences did not affect any results. Stimuli were created with the Psychophysics toolbox in Matlab (2017b, 2019a). One-dimensional receptive fields (RFs) were mapped along the azimuth (horizontal) or elevation (vertical) axes of the visual field using flashed bars. For azimuth maps, black or white vertical bars were presented one at a time at one of 17 evenly spaced azimuth locations from -37.8° to 115.8° (vertical meridian defined as 0°). Both the spacing between bar locations and the bar width was 9.6°. The bar height was 60° which spanned the entire vertical extent of the monitors. Bars were presented for 100 ms followed by 300 ms of gray screen. Bar location and luminance (black or white) were randomized. Each bar location and luminance were repeated for 20-30 trials.

Elevation (vertical) receptive field mapping (Fig. S2) was similar except horizontal bars at 6 evenly-spaced elevation (vertical) locations from -18.6° to 29.4° (horizontal meridian defined as 0°) were used. Both the spacing between bar locations and the bar height was 9.6°. The bar width was 155° which spanned the entire horizontal extent of the monitors.

### Habituation

Mice were habituated to head-fixation on the recording rig in daily sessions that increased progressively in duration from 10 minutes to >1 hour for at least 3 days before recording. To mimic recording conditions, head-fixed mice were positioned in a semi-enclosed plastic tube that prevented locomotion or large body movements^125^ while receptive field mapping stimuli were displayed. Mice were not water restricted or trained in any behavioral task. Sweetened condensed milk was given periodically in between runs of visual stimulation.

### Electrophysiological recordings

Mice were anesthetized under isoflurane (3% induction, 1-2% maintenance) and a small craniotomy was made (<0.5mm diameter for visual cortex; <1 mm along AP and <0.5mm ML for frontal cortex). The skull was thinned with a dental drill and the final layer of bone was removed with fine forceps and a 30g needle. The dura was left intact. Frontal cortex was targeted using stereotaxic coordinates (-0.5 to +0.5 mm AP, 0 to +0.5 mm ML from Bregma) and V1 and AM/PM were targeted using intrinsic signal imaging maps. Silicone elastomer (Kwikcast or a combination of Kwikcast and Dowsil 3840) were used to seal the craniotomy after the surgery and between recordings. Mice were allowed to recover for 4-24 hours before recording.

Mice were head-fixed in the recording rig and the silicone elastomer covering the craniotomy was removed and replaced with a bath of artificial cerebral spinal fluid (in mM: 150 NaCl, 4 KCl, 10 HEPES, 2 CaCl_2_, 1 MgCl_2_, pH 7.4, 310 mOsm). A Neuropixels 1.0 probe was inserted into the craniotomy using a micromanipulator at a 10-20º deviation from vertical (probe approached from the animal’s posterior side and was parallel to the midline). Probes were lowered at 2µm/s, travelling 2-3mm deep in frontal cortex and 1.3-1.8mm in visual cortex. Probes were coated with the red fluorescent lipophilic dye DiI prior to insertion. The ground and reference pads of the probe were connected to an Ag/AgCl wire in the ACSF bath. AP (30 kHz) and LFP (2.5 kHz) band data of the bottom 384 channels were acquired with SpikeGLX software in external reference mode. The probe was allowed to settle for 15-30 minutes prior to data acquisition. In some recordings, the ACSF bath was replaced with a drop of 1.5% agarose in ACSF (Type III-A, Sigma) covered in mineral oil. At the end of the recording, the probe was withdrawn and the craniotomy was rinsed with ACSF and covered with silicone elastomer and the animal returned to its home cage. Up to 6 recordings were performed from each craniotomy on separate days.

### Optogenetic silencing of visual cortex

Silencing experiments were performed in PV-ChR2 mice which express channelrhodopsin (ChR2) in parvalbumin-expressing cortical inhibitory interneurons. Optogenetic activation of cortical inhibitory neurons with blue light strongly silences spiking of neighboring excitatory neurons^48,74,126^ (Fig. S5). A fiber coupled laser diode (LP488-SF20G, Thorlabs) and galvanometer-based laser scanning system (GVS212 galvo, FTH254-1064 scan lens, PAF2P-A10A fiber collimator, Thorlabs) were used to deliver blue light to visual cortex sites targeted through the intact skull. The spot size (gaussian profile, full-width at half-maximum) at the brain surface was 36 µm. Binocular V1 (V1B), Monocular V1 (V1M), and the border between AM and PM (AM/PM) were targeted using previously acquired intrinsic signal imaging maps in each subject. We verified that the locations of our recordings in frontal cortex receive prominent projections from V1 and especially HVAs AM and PM by analyzing data from anterograde tracing experiments from the Allen Brain Connectivity Atlas ^55^(http://connectivity.brain-map.org/). We focused on AM/PM for inactivation and recording experiments based on these dense interconnections with frontal cortex.

For each recording, azimuth and elevation receptive fields were first mapped without any optogenetic manipulation (above, Methods, Visual stimuli) to identify neurons with RFs and their best bar locations (below, Methods, Analysis, RF Quantification). Then, azimuth RFs were mapped again during interleaved control trials and trials with optogenetic silencing (Fig. 4a). Visual stimuli were presented in trials of 12 sequential bar presentations (same timing and randomization as above) followed by a gray screen intertrial interval of 2.5 seconds between the end of the last bar and the start of the first bar of the next trial. On opto trials, the targeted site in visual cortex was illuminated with constant laser power (2-5 mW) for the entire duration of bar presentation (5 second duration; starting 0.35 seconds prior to the first bar presentation; ending 0.92 seconds after the last bar presentation). In some recordings, laser power was linearly ramped for 100ms at onset and offset. During interleaved control trials, bars were presented without laser illumination. There were 1-2 interleaved control trials per opto trial and a total of 20 opto trials and 20-40 control trials per opto site per recording. 2-3 opto sites were tested per recording.

#### Spike sorting

AP-band recordings were pre-processed and spike-sorted using Kilosort3^127^. Pre-processing involved band-pass filter (300-9000Hz), common median subtraction, motion correction, and whitening steps. Kilosort cluster output was manually curated in Phy (https://github.com/cortex-lab/phy). Clusters corresponding to high quality single units were manually identified based on neuron-like waveform, template amplitude distribution, and clear refractory period (<0.5% inter-spike intervals <1.3 ms in 99.9% of units). In rare cases, clusters were manually merged or split. Clusters were merged when they exhibited similar spike waveforms and clear refractory periods in their cross-correlograms. Splits were performed manually in the amplitude view when a cluster showed obvious outlier spikes. After manual curation, double-counted spikes, an artifact of template matching, were removed from each single unit by eliminating spikes occurring within 0.3 ms of a previous spike.

#### Definition of FS and RS units

Average spike waveform was used to distinguish regular spiking (RS) and fast spiking (FS) units. Spike waveforms of each unit were extracted from the pre-processed voltage trace at the unit’s peak channel. Individual waveforms were un-whitened and re-aligned to the peak (initial negative deflection) before averaging. Average waveform was normalized by dividing by the magnitude of the peak. The spike width was defined as the time interval between the peak and the maxima of the subsequent positive-deflecting trough. The trough amplitude was defined as the height of the trough. As in prior work ^70^, units with spike width < 0.5 ms and trough amplitude > 0.3 were classified as FS and the remaining units were classified as RS.

### Analysis

#### RF quantification

Spiking responses to RF mapping stimuli were binned (1ms) in peri-stimulus time histograms (PSTHs). A separate PSTH was calculated for each bar location and luminance (black or white). The PSTHs were smoothed in time with a symmetric 40 ms wide moving window average. Smoothed PSTHs were then z-scored using the mean and standard deviation of the 100 ms baseline period preceding stimulus onset across all bar locations and luminances. A spatial profile of the spiking response across bar locations was calculated as the baseline-subtracted average spike rate from 20 to 250 ms after stimulus onset, then fit with a 1D Gaussian (see below).

RFs were determined separately for black or white bars. Units were classified as having a spatial RF if bar responses of either color fulfilled 3 criteria:

1. the z-score from 20 to 200 ms after stimulus onset was >=5 z-scores for two adjacent bar locations
2. the largest peak value of the baseline-subtracted smoothed PSTH from 20 to 200 ms after stimulus onset across all bar locations was >=5 spikes/s
3. the Gaussian fit of the spatial profile had r^2^>0.5

For each black or white RF, the best bar location was defined as the location whose smoothed PSTH had the largest peak value from 20 to 200 ms after stimulus onset. For units with both black and white RFs, the best luminance was defined as the one whose best bar location had the largest peak value. When a unit had a receptive field for only one luminance, this luminance was considered the best luminance. Except for analysis of luminance selectivity (Fig. 2), receptive field analyses were conducted using each unit’s best luminance and the best bar location for that luminance.

To calculate peak latency, the PSTH for the best bar location was smoothed in time (20 ms symmetric moving window average) and the latency of the peak within 20 to 200 ms after stimulus onset was defined as the peak latency.

The spatial properties of the RF were derived by fitting the spatial profile of spiking responses to a 1D Gaussian of the form

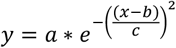

where the spiking response (y) was expressed as a function of bar location (x). Fit parameters were used to define the RF center location (b) and the RF width (full-width at half maximum,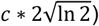.

#### Smoothing, normalization, and best-bar alignment for figures

All figures showing the time course of spiking including RF maps depict PSTHs smoothed in time with a 20 ms symmetric moving window average. For normalized population averages, the smoothed PSTH was baseline-subtracted (baseline firing rate calculated 0 to 100 ms before stimulus onset across all bar locations and luminance values) and divided by its maximal value before averaging across units. The spatial profile was divided by its maximal value before averaging across units. To align RF maps and spatial profiles to the best bar location, the data were shifted on the location axis (y-axis for RF maps and x-axis for spatial profiles) so that the best bar location occurred at 0º. After alignment, data values for bar locations that were not presented were not included in population averages.

#### Quantification of binocular bias

The maximal value of each frontal cortex unit’s RF spatial profile (see above) for binocular (<40º azimuth) and monocular (>40º azimuth) bar locations was used to compare responses to the binocular and monocular visual field.

#### Luminance (OFF/ON) selectivity

To quantify each frontal cortex RF unit’s OFF/ON selectivity, the responses to black and white bars were compared at a common set of bar locations that drove the strongest overall response in PSTHs that averaged across black and white luminances. The combined PSTHs for black and white bars were baseline-subtracted and smoothed with a 40 ms symmetric moving window average. The bar location with the greatest peak response in this combined black/white RF map was assigned and then PSTHs for this bar location and the two neighboring adjacent bar locations (best ±1 locations) were averaged separately for black and white bars. The peak value of the resulting black PSTH and white PSTH were used to calculate an OFF/ON selectivity index as follows

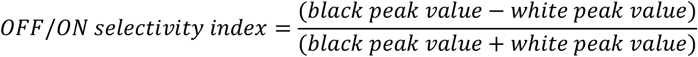

Units were divided into selectivity groups numbered from 1 to 5 (OFF/ON selectivity index ranges: [-1 to -0.5], [-0.5 to -0.2], [-0.2 to 0.2], [0.2 to 0.5], and [0.5 to 1]; Fig. 2C-D). In population averages of OFF and ON RF maps, each unit’s OFF and ON map were aligned to the combined best bar location (defined above) and normalized by the peak value across OFF and ON maps before averaging across units (Fig. 2D top). In population averages of OFF and ON responses (Fig. 2D, bottom), the PSTHs of the combined best bar location and the two neighboring adjacent bar locations were averaged together for each unit before normalizing to the peak and averaging across units.

#### Frontal cortex RF analysis during optogenetic silencing

The initial round of azimuth receptive field mapping was conducted prior to any optogenetic silencing was used to define frontal cortex units with RFs, and their best bar location and luminance. Subsequent analysis of responses during optogenetic silencing (“opto”) and interleaved trials without photostimulation (“control”) were restricted to these units and both RS and FS units were included.

RF maps and spatial profiles (at each unit’s best luminance) were constructed separately from opto trials and interleaved control trials, and sorted by the visual cortex site targeted for inactivation (Fig. 4B: Binocular V1 (V1B); monocular V1 (V1M); AM/PM). Data were baseline subtracted as previously described. Baselines were calculated separately for opto and control trials. Population averages of RF maps and spatial profiles were aligned to the best bar location as previously described. The best bar location was determined using the initial round of RF mapping data prior to optogenetic silencing rather than interleaved control trials during photostimulation. The response at this location was not significantly different between the initial versus interleaved control trials. Data were normalized by dividing by the peak value across both opto and control data before averaging across units. Response magnitude in opto and control trials (Fig. 4D) corresponds to the value of the spatial profile at the best bar location. Magnitude of opto reduction (Fig. 4E) was computed as the percent reduction of this value in opto trials relative to control trials. Histology

#### Perfusion and imaging

After the final recording, mice were injected with a lethal dose of phenotoin/pentobarbatol (Euthasol, IP) and transcardially perfused with 4% paraformaldehyde (PFA) in 1x PBS. The brain was extracted and kept in PFA for 24 hours and transfer to 1x PBS. In a subset of mice with multiple frontal cortex recordings, 100 μm coronal sections of frontal cortex were made on a vibratome (Leica VT1000S), DAPI stained, and mounted on glass slides (Vectashield H-1000-10). Red (DiI), green (autofluorescence), and blue (DAPI) fluorescent images of each slice were acquired at 4x magnification (BioTek Cytation 7).

#### Registration to Allen Brain Atlas

Images were aligned to the Allen Brain Atlas Common Coordinate Framework (CCF)^73^ (https://atlas.brain-map.org/) using ABBA software^128^. A series of automated registration steps were performed (2 rounds of DeepSlice registration, affine, and spline) followed by manual adjustment of the registration around the frontal cortex. DiI tracks corresponding to each recording were annotated in QuPath software, converted to CCF coordinates, and fit with a linear function. The CCF coordinate of each unit was obtained by first estimating each unit’s distance along the probe from the brain surface. This was defined as the distance between each unit’s peak channel (the channel with the largest waveform amplitude) and the channel corresponding to the brain surface (where the LFP signals diminish). The CCF coordinate of each unit was then defined as the CCF coordinate corresponding to this distance along the linear fit of the probe track. The CCF area (secondary motor cortex: MOs; dorsal and ventral anterior cingulate cortex: dACA, vACA) for each unit was obtained using annotation volume masks. Area boundaries in the saggital plane (Figs. 3C and S2C) correspond to linear fits to the coordinates of the most superficial unit in that area for all recordings. The surface boundary was estimated in a similar way using the coordinates of the surface channel for all recordings. The anterior/posterior CCF coordinate of Bregma was estimated to be 5.4 mm, and this was subtracted from unit CCF coordinates to obtain their anterior/posterior position relative to Bregma (Figs. 3C and S2C). The dorsal/ventral axis was also offset by 1mm to approximately place the surface at 0 mm.

---

**Figure S1.**
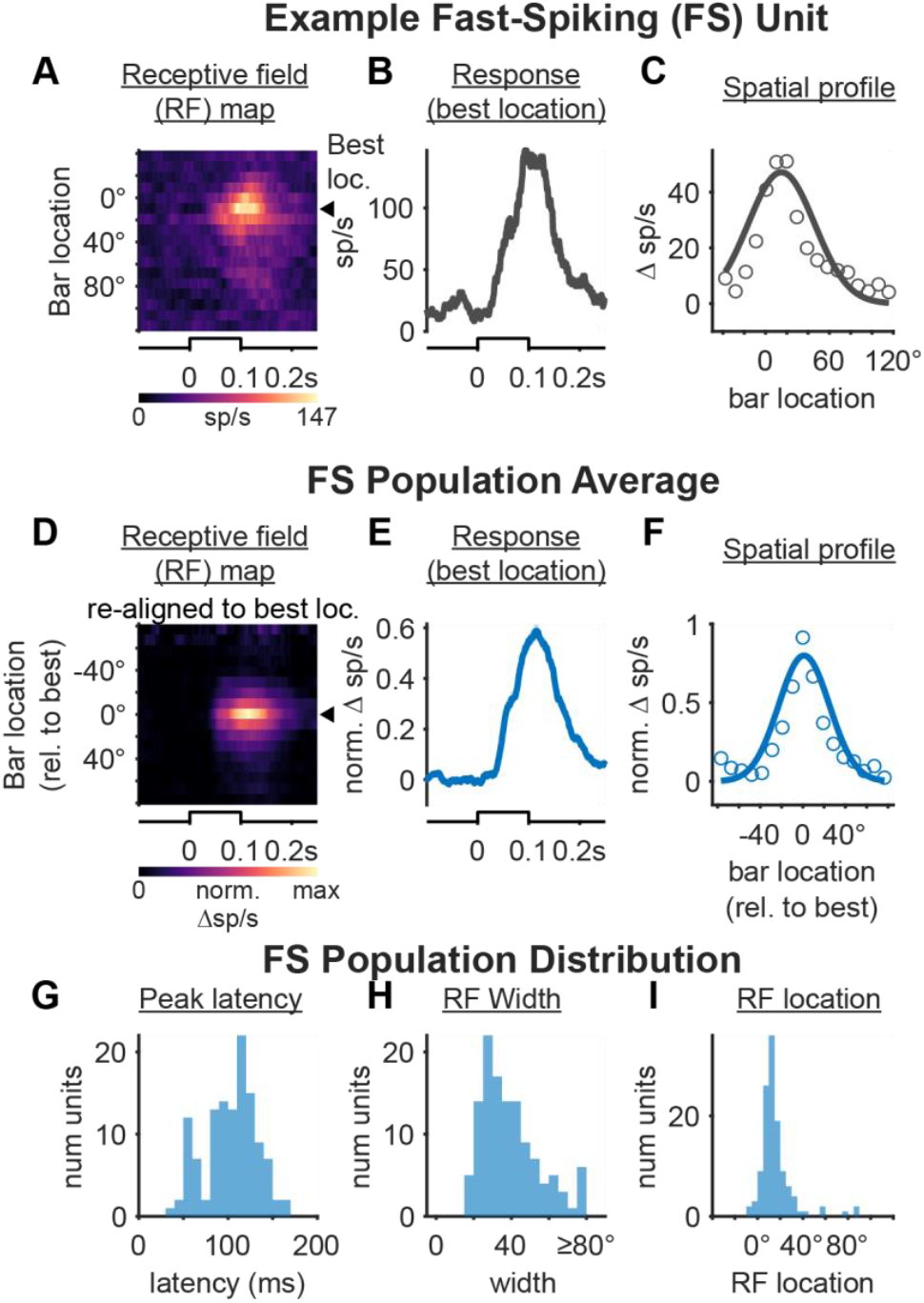
Visual receptive fields of frontal cortex fast spiking (FS) neurons. A. Receptive field map of an example frontal cortex fast-spiking (FS) neuron. Triangle indicates best bar location. Only responses to white bars shown. *B*. Same neuron as A, showing time course of best location response. C. Same neuron as A, showing spatial profile of RF responses (circles) with Gaussian fit (solid). D. Population RF map computed after aligning each neuron’s best bar response (color and location). (n=121 FS units, 47 recordings, 14 mice in D-I). E. Population average spiking response to the best bar location. F. Population average spatial profile (circles) with Gaussian fit (solid). G. Peak latency distribution calculated from each unit’s best bar response (109±19 ms, median +/-M.A.D.). H. Distribution of RF width calculated from each unit’s spatial profile. (36±9º, median +/-M.A.D. significantly different from RS *p* = 6.97e-7, rank sum test). I. Distribution of RF center locations (13±5º, median +/-M.A.D.).

**Figure S2.**
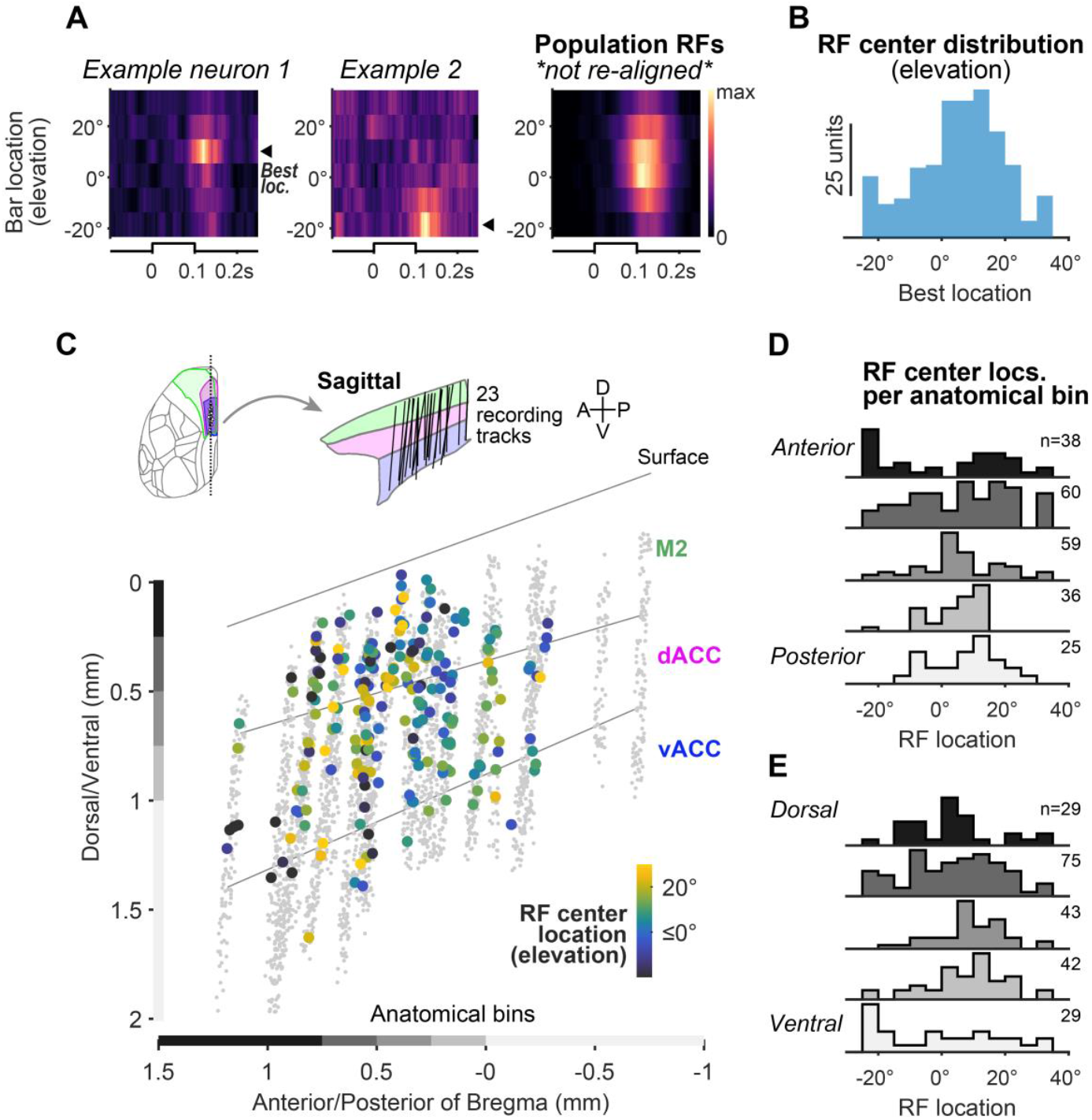
Elevation receptive fields in frontal cortex. A. *Left and middle*, Elevation receptive field maps of two example RS frontal cortex neurons mapped using horizontal bars. Triangles (best bar location) denote RF center. *Right*, Population average RF map (without re-alignment of best bar location; n=335 RS units, 47 recordings, 14 mice). *B*. Distribution of RF center locations in elevation, same neurons as A. Population average elevation receptive field map (computed without re-alignment of best bar location). (n=335 RS units, 47 recordings, 14 mice in B-C). C. *Top*, all recording tracks with fluorescent label (n=23), projected in the sagittal plane. *Bottom*, neurons with significant visual elevation RFs color-coded by center location (n=218 RS neurons), all other non-RF neurons in gray (n=3370 RS neurons). Lines denote area boundaries. Gray and black bars above axes indicate anatomical bins used in D and E (n=23 recordings, 5 mice; same units for C-E). For display purposes only, neuron coordinates were randomly jittered by ≤25 μm in each dimension. D. Distribution of RF center locations across the anterior/posterior axis using the 5 anatomical bins depicted in C. No correlation between RF location and anterior/posterior coordinate (r=0.005, p=0.94; Spearman correlation). E. Same as D for the dorsal/ventral axis. No correlation between RF location and dorsal/ventral (r = 0.07, *p* = 0.31; Spearman correlation) or medial/lateral coordinate (r = 0.008, *p* = 0.90; Spearman correlation)

**Figure S3.**
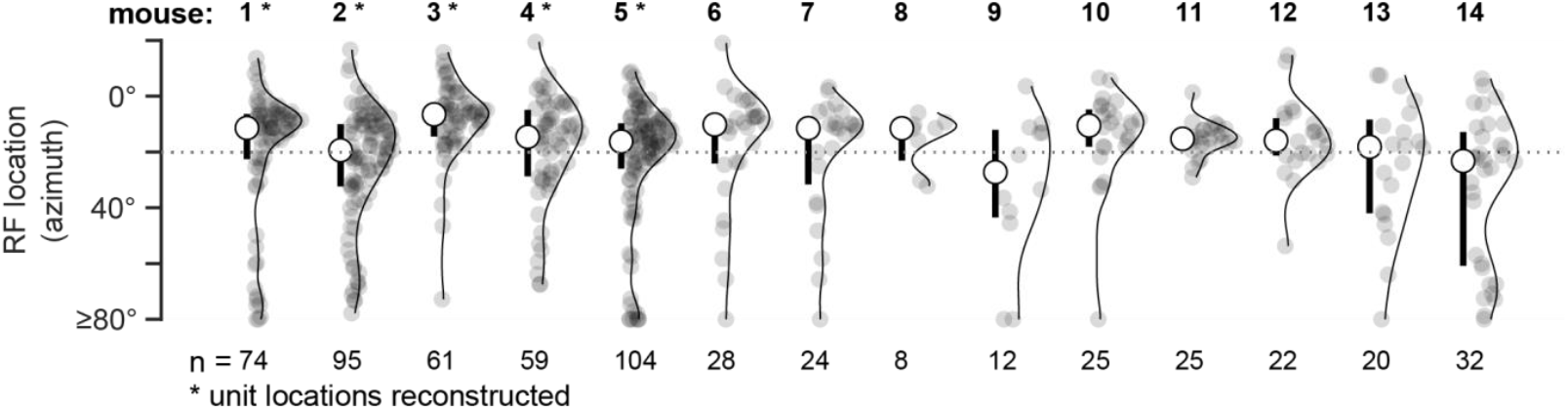
Frontal RF locations across mice. Distributions of frontal RS neuron azimuth RF locations for each mouse with markers denoting median and IQR. Asterisks indicate mice where recording sites were reconstructed with histology (Fig. 3 and Fig. S2). Dotted line marks 20º azimuth; binocular region spans central ±20°.

**Figure S4.**
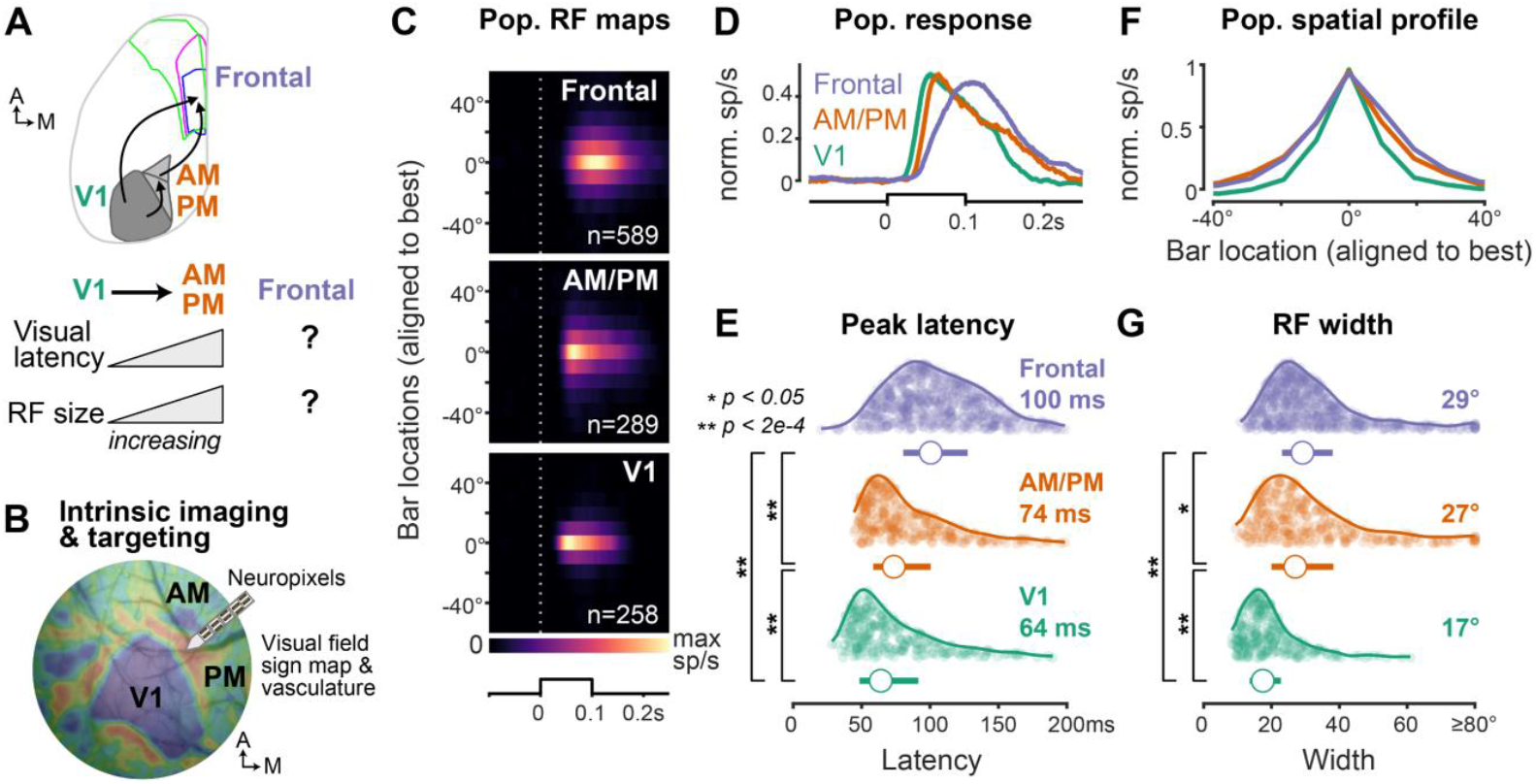
Comparison of RF properties in V1, higher visual areas, and frontal cortex. A. Schematic showing known anatomical connections from primary visual (V1) and antero- or posteromedial (AM/PM) higher visual areas to frontal cortex (See Methods for verification from Allen Brain Connectivity Atlas). *B*. Intrinsic signal imaging visual field sign map showing visual cortical area boundaries relative to vasculature overlay. C. Population RF maps for frontal, AM/PM, and V1 RS neurons (aligned to best bar location, defined as 0º; frontal: n=589, 47 recordings, 14 mice; AM/PM: n=289, 17 recordings, 6 mice; V1: n=258, 15 recordings, 5 mice). D. Population average spiking response at best bar location for each area. Mean ± SEM plotted. E. Distribution of peak latency for each area with median ± IQR below. All groups significantly different from one another (V1: 64 ± 43; AM/PM: 74 ± 42; Frontal: 101 ± 47, *p*=5.7e-43, Kruskal-Wallis, post hoc Dunn’s test). F. Population average spatial profile for each area (normalized to peak at best location). Mean ± SEM plotted. G. Distribution of RF width (defined as full-width at half-max of best-fit Gaussian) for each area with median ± IQR below. All groups significantly different from one another V1: 17.4 ± 9.2; AM/PM: 27.0 ± 18.3; Frontal: 29.1 ± 15.0, *p*=5e-48 Kruskal-Wallis, post hoc Dunn’s test).

**Figure S5.**
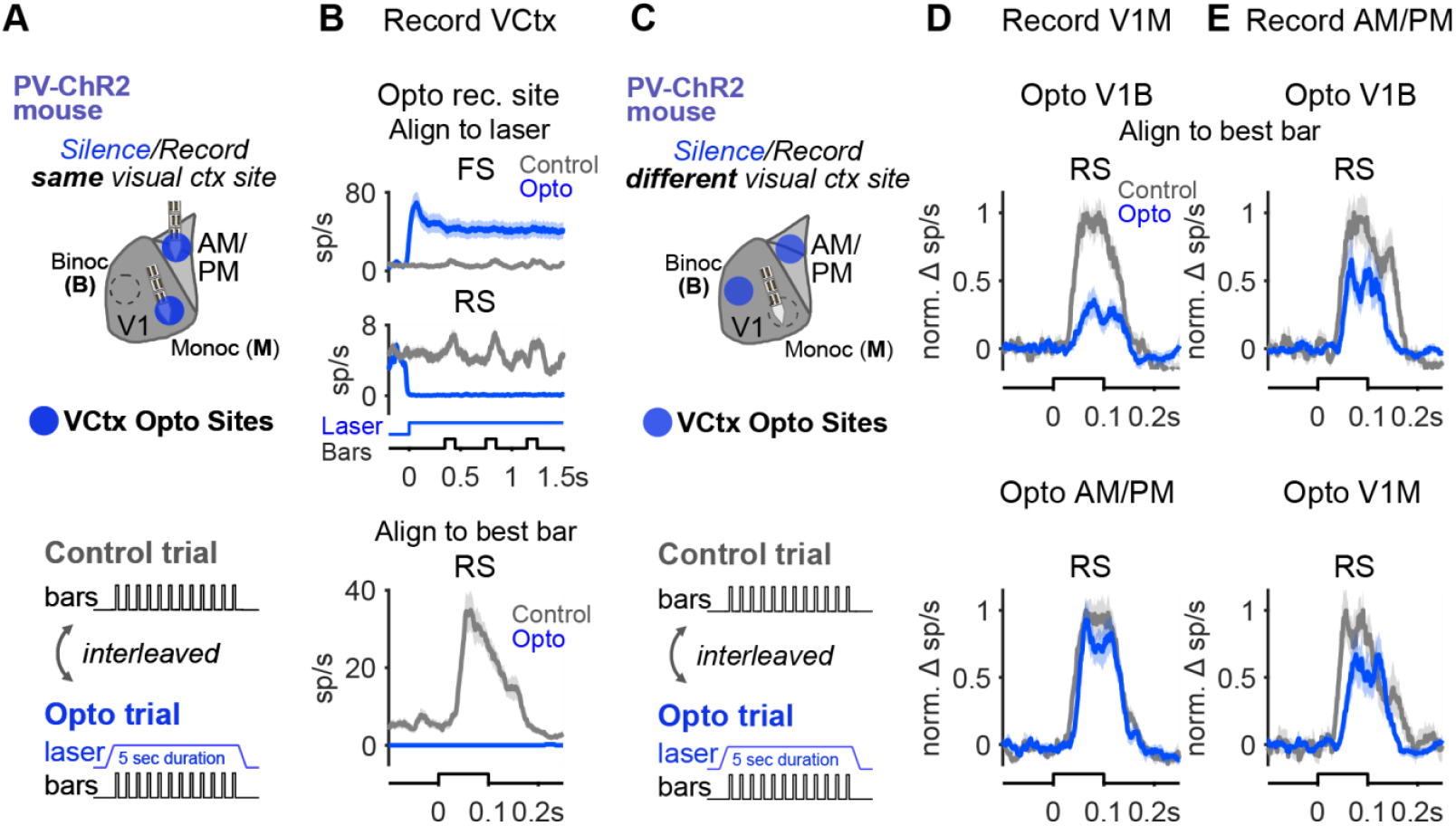
Optogenetic silencing of visual cortex. A. Schematic of experiment in B to assess cortical silencing at the targeted optogenetic site in visual cortex. Recordings were performed in V1M and AM/PM and the laser was targeted to the same location as the recording site. Vertical bars for receptive field mapping were presented during opto and control trials *B*. Average response of fast-spiking (FS) and regular-spiking (RS) neurons during control trials (gray) and opto trials (blue) when the laser was targeted to the recording site. Neurons from V1M and AM/PM recordings were pooled (n=113 RS neurons and 36 FS neurons, 2 recordings, 1 mouse). Top panels, aligned to laser. Bottom panel, RS neurons with RFs aligned to onset time of each neuron’s best bar (n=42 neurons). Note strong laser activation of putative ChR2-expressing PV+ FS interneurons and silencing of putative RS excitatory neurons. C. Schematic of experiments in D and E to assess effects of visual cortical silencing on distal visual cortical sites. D. Normalized average spiking response of V1M RS neurons during opto over V1B (top) or AM/PM (bottom). Responses are aligned to the onset of each unit’s best bar. Control trials in gray and opto trials in blue. n=26 neurons, 1 recording, 1 mouse. E. Same as D for AM/PM RS neurons during opto over V1B (top) or V1M (bottom). N=16 neurons, 1 recording, 1 mouse.

